# *Tet2* negatively regulates memory fidelity

**DOI:** 10.1101/843581

**Authors:** Kristine E. Zengeler, Caroline P. Gettens, Hannah C. Smith, Mallory M. Caron, Xinyuan Zhang, Alexandra H. Howard, Andrea R. Boitnott, Alex R. Gogliettino, Anas Reda, Beth G. Malachowsky, Chun Zhong, Hongjun Song, Garrett A. Kaas, Andrew J. Kennedy

## Abstract

Despite being fully differentiated, DNA methylation is dynamically regulated in post-mitotic glutamatergic neurons in the CA1 of the hippocampus through competing active DNA methylation and de-methylation, a process that regulates neuronal plasticity. Active DNA methylation after learning is necessary for long-term memory formation, and active DNA de-methylation by the TET enzymes has been implicated as a counter-regulator of that biochemical process. We demonstrate that *Tet2* functions in the CA1 as a negative regulator of long-term memory, whereby its knockdown across the CA1 or haploinsufficiency in glutamatergic neurons enhances the fidelity of hippocampal-dependent spatial and associative memory. Loci of altered DNA methylation were then determined using whole genome bisulfite sequencing from glutamatergic *Tet2* haploinsufficient CA1 tissue, which revealed hypermethylation in the promoters of genes known to be transcriptionally regulated after experiential learning. This study demonstrates a link between *Tet2* activity at genes important for memory formation in CA1 glutamatergic neurons and memory fidelity.

## Introduction

Active DNA methylation in neural tissue is necessary for the formation and maintenance of long-term memory (Day and Sweatt, 2010; Levenson et al., 2006; Miller et al., 2010). DNA methyltransferases (DNMTs) catalyze the transfer of a methyl group from S-adenosyl methionine (SAM) to the 5′ position of cytosine (Jaenisch and Bird, 2003; Morris and Monteggia, 2014). 5-methylcytosine (5mC) is a highly stable epigenetic mark that is known to alter the transcriptional rate of genes, being associated with transcriptional silencing or activation when present in the promoter or body of genes (Bintu et al., 2016; Yang et al., 2014). Whether or not DNA methylation in promoters blocks gene expression is dependent upon the structure and binding modality of different transcription factors (Yin et al., 2017). Experiential learning induces active DNA methylation at genes that have been identified to be necessary for long-term memory formation in circuits that encode spatial, associative, and reward memories (Day et al., 2013; Duke et al., 2017; Guo et al., 2011). The changes to the DNA methylome induced in neural tissue after learning are dynamically regulated (Baker-Andresen et al., 2013; Lister and Mukamel, 2015), and 5mC can be oxidatively converted to 5-hydroxymethylcytosine (5hmC) by the ten-eleven translocation (TET) enzymes and removed through base excision repair, in effect causing the erasure of the DNA methylation mark (Pastor et al., 2013; Rasmussen and Helin, 2016; Tahiliani et al., 2009). Such dynamism paired with the essential role that DNA methylation plays in learning and memory suggests that active DNA de-methylation, mediated by the TET enzymes, may be a target for therapeutic intervention to affect memory function (Kennedy and Sweatt, 2016).

There are three TET enzyme genes (*Tet1-3*) in mammals, all of which produce proteins that can catalyze the oxidation of 5mC (He et al., 2011; Ito et al., 2011). *Tet1* has been implicated in the regulation of glutamate receptor trafficking and synaptic scaling (Meadows et al., 2015; Sweatt, 2016; Yu et al., 2015) and as a negative regulator of memory function, where *Tet1* gene deletion and silencing enhances long-term memory and inhibits memory extinction (Kumar et al., 2015; Rudenko et al., 2013), while *Tet1* upregulation impairs long-term memory (Kaas et al., 2013). *Tet3* expression regulates synaptic scaling and glutamatergic synaptic transmission, and is upregulated in the hippocampus after memory retrieval induces engram destabilization (Liu et al., 2018; Yu et al., 2015). However, it has in general been difficult to relate these manipulations of *Tet1* and *Tet3* activity in neural tissue to precise alterations in DNA methylation patterns at genes associated with memory function, something that has stymied a clearer understanding for how dynamic DNA methylation facilitates memory consolidation and recall.

Despite being highly expressed in the adult CNS, little is known about how *Tet2* regulates DNA methylation in post-mitotic neurons, its genomic targets in neural tissue, or whether it facilitates memory function (Antunes et al., 2019). Recently, it was demonstrated that *Tet2* drives adult neurogenesis in the dentate gyrus in mice, and that *Tet2* knockout in neural progenitor cells (NPCs) impairs cognition (Gontier et al., 2018). *Tet2* was also found to be elevated in the brain tissue of humans with Alzheimer’s Disease and a mouse model of the disease (Carrillo-Jimenez et al., 2019). Here, we test the idea that *Tet2* can be targeted in post-mitotic glutamatergic neurons to enhance the fidelity of long-term spatial memory by functionally increasing DNA methylation. Additionally, we determine the genomic regions of hypermethylation that *Tet2* reduction induces in the CA1 region of the hippocampus.

## Results

### *Tet2* knockdown in the CA1 of the hippocampus enhances object location memory

To test the hypothesis that active DNA de-methylation mediated by *Tet2* in the CA1 of the hippocampus impairs long-term spatial memory, an AAV engineered to express shRNA that targets *Tet2* transcripts was surgically injected into the dorsal CA1 in both hemispheres of adult C57B6/J mice (Figure 1a), controlled with an AAV that expresses a scramble shRNA injected into littermates as previously described (Kaas et al., 2013). Two weeks post-surgery, a group of these mice were sacrificed to demonstrate that *Tet2*-shRNA elicited 41% *Tet2* knockdown in the CA1 relative to scramble shRNA (Figure 1b). No differences in baseline activity or anxiety were observed in mice injected with *Tet2*-shRNA compared to scramble, as measured by open field assessment (Figure S1). *Tet2* deficient mice were also submitted to the object location memory (OLM) paradigm two weeks post-surgery (Figure 1c). Briefly, OLM begins by habituating mice for several days to a box that contains a spatial cue on one wall, before being exposed for 10 m to two identical objects (50 mL beakers) equidistance from the spatial cue. To test for 1 d long-term memory, each mouse reenters the box where one object remains in its familiar location and the other object has been moved to a novel location, distal to the spatial cue. Novelty drives increased interaction with the object in the novel location and memory is inferred by a discrimination index (DI) between the interaction times of object in the novel and familiar locations. *Tet*2-deficient mice demonstrated a significant enhancement in discrimination of the object in the novel location versus the familiar (Figure 1d). We interpret this result to mean that *Tet2* activity in the CA1 of the hippocampus inhibits long-term spatial memory function. Nevertheless, viral-mediated *Tet2* knockdown throughout the dorsal CA1 fails to identify which cell types govern this memory enhancement. Since *Tet1* and *Tet3* had been implicated in glutamate signaling, we hypothesized that *Tet2* activity in post-mitotic glutamatergic neurons in the CA1 undermines long-term memory function.

**Figure 1.**
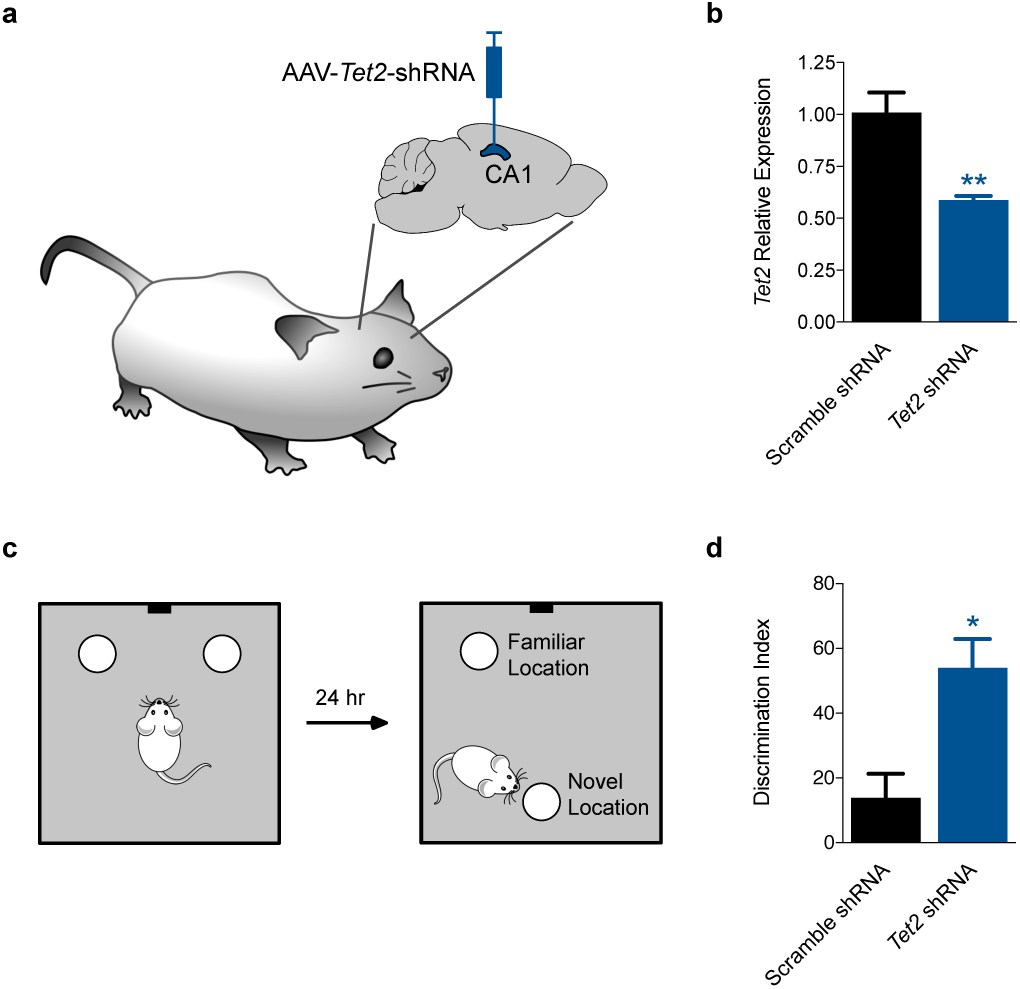
Tet2 knockdown in the CA1 of the hippocampus enhances object location memory. (A) AAVs engineered to expressed either *Tet2*-targeted or scramble shRNAs were injected into the CA1 region of the hippocampus in mice. (B) *Tet2*-shRNA mediated mRNA knockdown as determined by qRT-PCR in CA1 tissue 14 days post-surgery (N = 3-5 per group). (C) 24 hr object location memory paradigm design. (D) *Tet2*-deficient mice demonstrate enhanced discrimination in 24 hr object location memory (N = 7 per group) Data shown are represented as mean ± sem. Two-tailed Student’s t test. *p < 0.05 and **p < 0.01.

### The conditional knockout of *Tet2* in glutamatergic neurons enhances long-term spatial memory

To test the idea that this enhanced memory phenotype is facilitated by *Tet2* activity in post-mitotic glutamatergic neurons, the conditional knockout of *Tet2* selectively in these cells was facilitated using mice containing a *Tet2* floxed allele (Moran-Crusio et al., 2011) and Camk2a-driven Cre expression, which is well characterized to express active Cre recombinase in glutamatergic neurons selectively in the CA1 and forebrain, with recombination occurring during the third week after birth (Tsien et al., 1996). In order to match our level of *Tet2* knockdown observed in the previous experiment, heterozygous *Tet2* +/flox Camk2a-Cre mice were used along with *Tet2* +/flox controls to elicit 50% knockout in glutamatergic neurons (Figure 2a). These conditional knockout mice did not have any apparent alterations in activity or anxiety as measured by open field assessment (Figure S2), and mice were then submitted to the OLM paradigm for either short-term (2 h) or long-term (1 d or 7 d) memory test (Figures 2b and S2). OLM elicits a transient memory that is thought to last no more than a few days (Ciernia and Wood, 2014). Similar to *Tet2* knockdown mice, *Tet2* +/flox Camk2a-Cre mice demonstrated significantly enhanced 1 d OLM, and further showed a significant OLM phenotype at 7 d at which point *Tet2* +/flox controls fail to discriminate (Figure 2c). Representative traces during the 7 d test show a *Tet2* +/flox mouse spending equal time exploring both beakers, while a *Tet2* +/flox Camk2a-Cre mouse discriminates toward the object in the novel location (Figure 2d). Notably, short-term 2 h OLM was unaffected in *Tet2* knockout mice (Figure S2).

**Figure 2.**
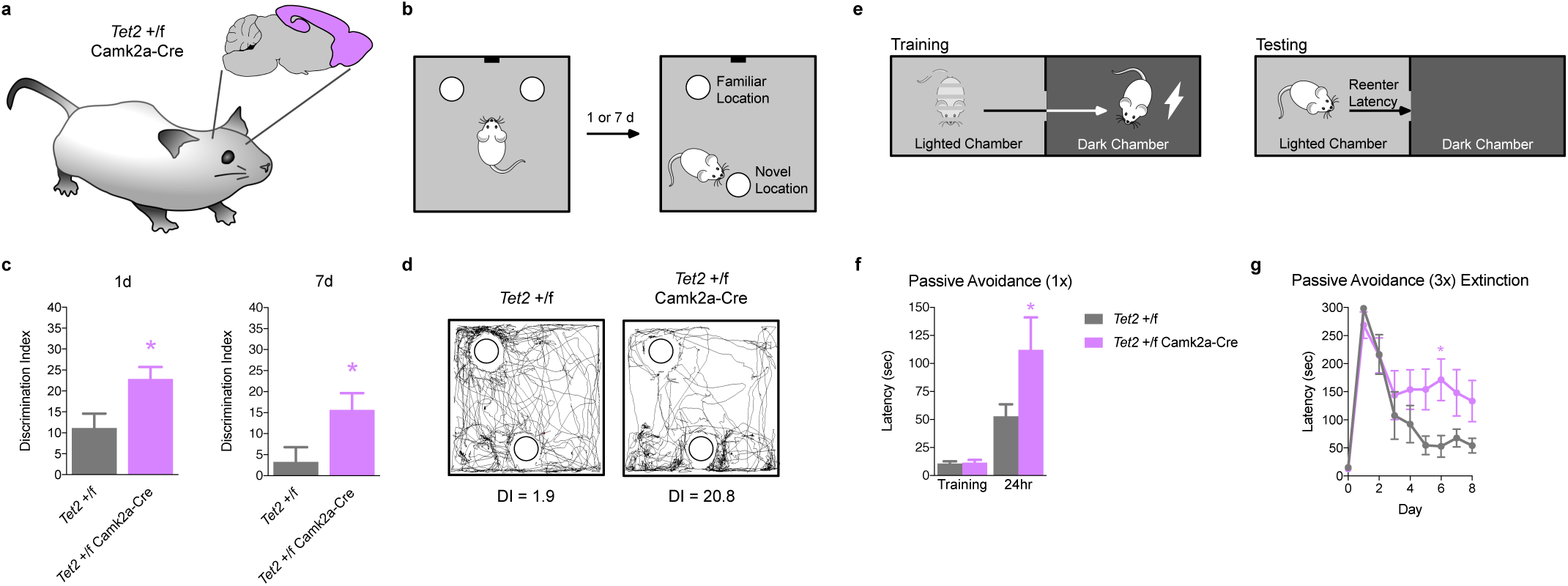
*Tet2* knockout in glutamatergic neurons enhances long-term spatial memory. (A) *Tet2* +/flox Camk2a-Cre mice were used along with *Tet2* +/flox controls to elicit 50% knockout post-developmentally in glutamatergic neurons in the CA1 and forebrain. (B) OLM was tested for either long-term 1 d or remote 7 d memory. (C) *Tet2* +/flox Camk2a-Cre mice have enhanced 1 d and 7 d OLM. *Tet2* +/flox controls failed to demonstrate significant memory at the remote 7 d timepoint (N = 9-18 per group). (D) Representative traces of *Tet2* +/flox Camk2a-Cre and *Tet2* +/flox mice during the remote 7 d test. (E) Passive avoidance memory paradigm design. (F) *Tet2* +/flox Camk2a-Cre mice have increased latencies to reenter the dark chamber 1 d after weak (1x) training (N = 10-17 per group). (G) *Tet2* +/flox Camk2a-Cre mice show a deficits in memory extinction in the passive avoidance paradigm over subsequent testing sessions following strong (3x) training (N = 10 per group). Data shown are represented as mean ± sem. Two-tailed Student’s t test or two-way ANOVA with Sidak’s multiple comparisons test. *p < 0.05 and **p < 0.01.

To determine if this enhancement applied to contextual associative memory, *Tet2* +/flox Camk2a-Cre mice were conditioned using the passive avoidance paradigm (Figure 2e). Mice were placed into one chamber of a shuttle box that was lighted and allowed to pass through a doorway into the other chamber of the box that had been kept dark. Upon entering the dark chamber, mice were trained to associate that context with a paired 2 s 0.5 mA foot shock. Memory of the association was tested 1 d later when the mice were returned to the lighted chamber, whereby all groups demonstrated an increased latency to reenter the dark chamber. When mice were administered weak training, in the form of a single foot shock pairing, *Tet2* +/flox Camk2a-Cre mice show a significantly increased latency to reenter the dark chamber relative to control Tet2 +/flox mice (Figure 2f). We interpret this increased latency to reenter the dark chamber 1 d after weak training to be representative of a stronger association between the dark chamber and the threat of being startled.

A previous study characterized a memory extinction deficit in *Tet1* knockout mice (Rudenko et al., 2013). To test whether memory extinction is also impaired in *Tet2* +/flox Camk2a-Cre mice, a strong training protocol for the passive avoidance paradigm was used, in the form of three separate entries into the dark chamber over a 15 m training window, each accompanied by a 2 s 0.5 mA foot shock. Following this strong training protocol, both *Tet2* knockout and control mice demonstrate profound latencies to reenter the dark chamber over a 300 s testing session (Figure 2g). Mice were then retested subsequently each day for a total of 8 d. Control mice demonstrate archetypal memory extinction behavior, where latencies to reenter the dark chamber are significantly reduced over several testing days. *Tet2* +/flox Camk2a-Cre mice show a significant deficit in memory extinction as compared to controls. Extinction involves processing new information about the dark chamber that inhibits the expression of the previously acquired memory associating the dark chamber with a threat. *Tet2* +/flox Camk2a-Cre mice demonstrated enhanced long-term memory after weak training and a reluctance to extinguish a strong threat memory over subsequent retests (Figure 2f,g). We interpret these results to indicate that the reduction of *Tet2* in glutamatergic neurons in the CA1 and forebrain enhances the fidelity of the original memory phenotype and inhibits attempts to attenuate that memory, rather than an alternative explanation that the mice have enhanced cognition, more broadly.

### The conditional knockout of *Tet2* in glutamatergic neurons of the CA1 results in the hypermethylation of genes associated with memory formation

To test whether *Tet2* activity in CA1 glutamatergic neurons acts to reduce DNA CpG methylation, whole genome bisulfite sequencing (WGBS) of *Tet2*-deficient CA1 tissue was performed. The CA1 regions of *Tet2* +/flox Camk2a-Cre and *Tet2* +/flox mice were dissected, DNA extracted, fragmented, and bisulfite converted prior to Illumina sequencing to an average depth of 30x coverage across all CpG sites in the mouse genome per group (N = 6 mice per group). Bisulfite converted reads were then filtered for quality, mapped to the mouse genome (mm10), and the methylation status for each CpG was extracted (Bismark). The mouse genome was divided into windows, each containing 25 CpGs, and differential methylation was determined using the EdgeR filter, which can be used to detect differences in the ratio of methylated to unmethylated reads within a given window (FDR < 0.05).

A large majority of DNA methylation in mammalian tissue occurs in a CpG context. While no significant global difference in cytosine CpG methylation was detected between *Tet2* +/flox Camk2a-Cre and *Tet2* +/flox CA1 tissue (Figure 3a), 893 windows were found to be differentially methylated regions (DMRs), and these *Tet2* +/flox Camk2a-Cre DMRs were mostly hypermethylated (Figure 3b). When compared to the CpG methylation of all windows across the genome, these DMRs contain on average significantly less methylation than the rest of the genome. A cumulative distribution of the percent methylation of all windows across the genome as compared with the percent methylation of *Tet2* +/flox Camk2a-Cre and *Tet2* +/flox DNA at the 893 DMRs revealed that about half of the DMRs occur at regions of the genome with less than 50% CpG methylation, and there is a significant shift toward higher methylation for *Tet2* +/flox Camk2a-Cre relative to *Tet2* +/flox samples (Figure 3c). This indicates that these DMRs are not randomly distributed throughout the genome, and it was found they are significantly associated with gene bodies and even more highly concentrated at gene promoters (Figure 3d). To test whether these DMRs were preferentially associated with genes that are transcriptionally regulated by experiential learning, we compared their relative abundance to genes with altered transcription in the CA1 of the hippocampus 1 h after contextual fear conditioning (Kennedy et al., 2016). *Tet2* +/flox Camk2a-Cre DMRs were even more highly associated (> 4-fold more likely than random) with the promoters of these experience-regulated genes (ERGs) (Figure 3d).

**Figure 3.**
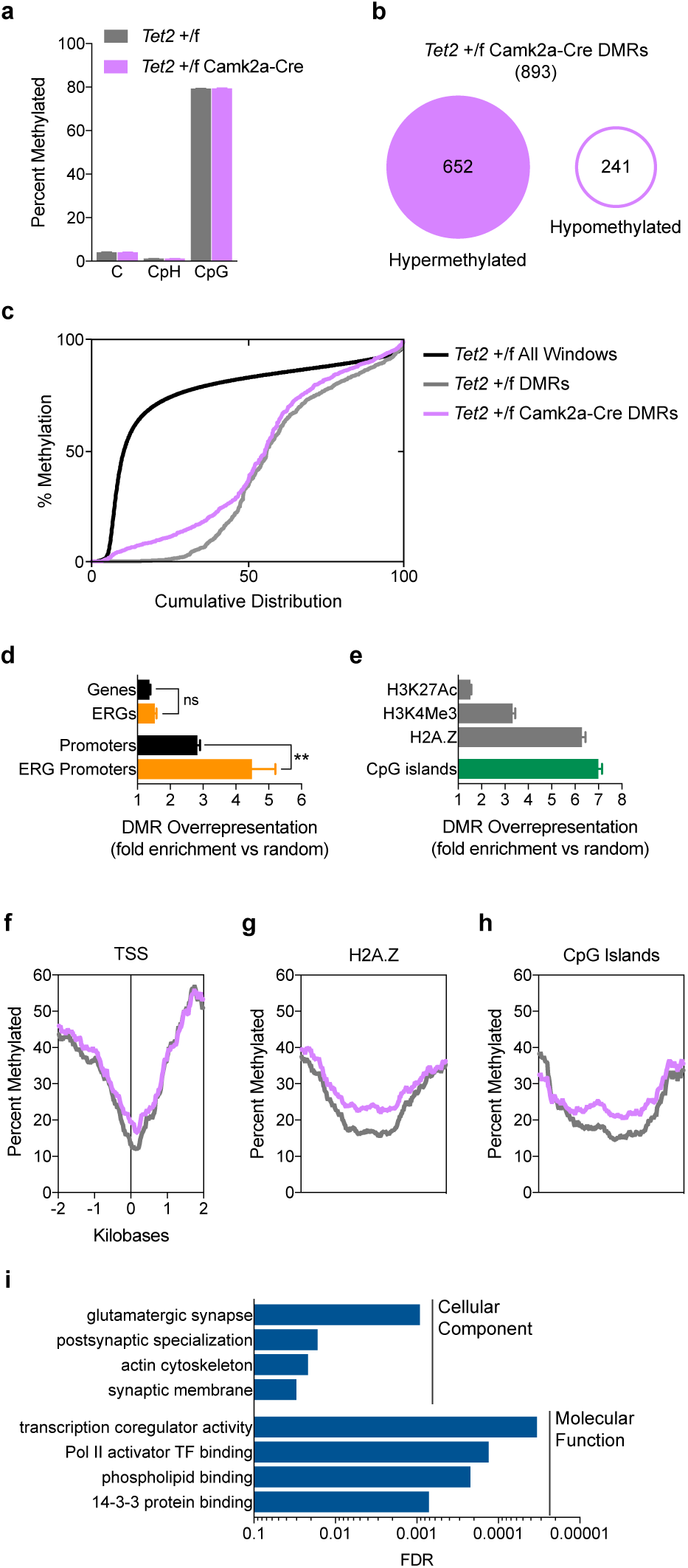
Whole genome bisulfite sequencing of CA1 tissue from *Tet2* knockout mice. (A) Global changes in cytosine methylation between *Tet2* +/flox Camk2a-Cre and *Tet2* +/flox CA1 tissue were not observed. (B) There were 893 *Tet2* +/flox Camk2a-Cre DMRs identified, most being hypermethylated. (C) *Tet2* +/flox Camk2a-Cre DMRs were associated with relatively unmethylated regions of the genome and were significantly hypermethylated relative *Tet2* +/flox tissue. KS test, P < 0.001. (D) Overrepresentation of DMRs at genomic features and ERG promoters. (E) Overrepresentation of DMRs at regions associated with H3K27Ac, H3K4Me3, and H2A.Z occupancy, as well as CpG islands (eo > 0.6). (F) Average percent methylation of DMRs +/-2kb from the TSSs. *Tet2* +/flox Camk2a-Cre marked as lavender and *Tet2* +/flox as grey. (G) Average percent methylation of DMRs at H2A.Z binding sites. *Tet2* +/flox Camk2a-Cre marked as lavender and *Tet2* +/flox as grey. (H) Average percent methylation of DMRs at CpG islands. *Tet2* +/flox Camk2a-Cre marked as lavender and *Tet2* +/flox as grey. (I) The gene ontology of genes that were associated with DMRs. Data shown are represented as mean ± sem. EdgeR filter for differential methylation in WGBS (FDR < 0.05) or Two-way ANOVA with Sidak’s multiple comparisons test. *p < 0.05.

To determine whether the DMRs were associated with other epigenetic marks found at activate promoters, the relative distribution of DMRs was also correlated with H3K27Ac and H3K4Me3 ChIPseq datasets collected from the mouse hippocampus (Gjoneska et al., 2015). The acetylation of H3K27 is associated with both active enhancers and promoters, while the trimethylation of H3K4 is highly correlated with active promoters. DMRs are overrepresented at both histone modifications, but more so for H3K4Me3, indicating that changes in DNA methylation associated with *Tet2* +/flox Camk2a-Cre preferentially target active gene promoters (Figure 3e). An analysis of the percent methylation of DMRs that reside +/-2kb around transcriptional start sites (TSSs) indicated increased methylation across promoters and especially around TSSs (Figure 3f). DMRs were even more associated with the histone variant H2A.Z (Figure 3g), which is found in the +1 and −1 nucleosomes at transcriptional start sites in the hippocampus and is ejected at ERGs after experiential learning (Stefanelli et al., 2018). Unsurprisingly, given the tight association of H2A.Z and unmethylated CpG islands (oe >0.6), DMRs were also concentrated at CpG islands (Figure 3h), with 77% of the DMRs being hypermethylated. Lastly, genes that were associated with DMRs were grouped by gene ontology, using the cellular component and molecular function categorizations available through WebGestalt. Gene categories that were significantly targeted by the *Tet2* knockout in the CA1 were those that encode for proteins associated with glutamatergic synapses and regulators of gene transcription (Figure 3i).

Many of the gene loci containing *Tet2* +/flox Camk2a-Cre DMRs encode for regulators of transcription that are activated shortly after learning, including the genes for *Nr4a3, Egr1*, and *Nedd9* (Figure 4a). RNA sequenced from the CA1 region of the hippocampus of mice 1 h after contextual fear conditioning, compared with naïve mice, showed significantly increased read density after learning at these three genes (Kennedy et al., 2016). Since these genes are actively expressed in hippocampal tissue, it follows that there is almost no DNA methylation around their TSSs (Figure 4b). Nucleosomes surrounding active TSSs often contain the H2A.Z histone variant, and of the 893 DMR identified due to *Tet2* knockout, 481 overlap with H2A.Z binding sites identified in the hippocampus (Stefanelli et al., 2018). WGBS allows for the interrogation of DNA methylation with single cytosine resolution, and we found that changes in methylation across these DMRs can occur broadly or at a subset of CpG sites within the 25 CpG window (Figure 4c).

**Figure 4.**
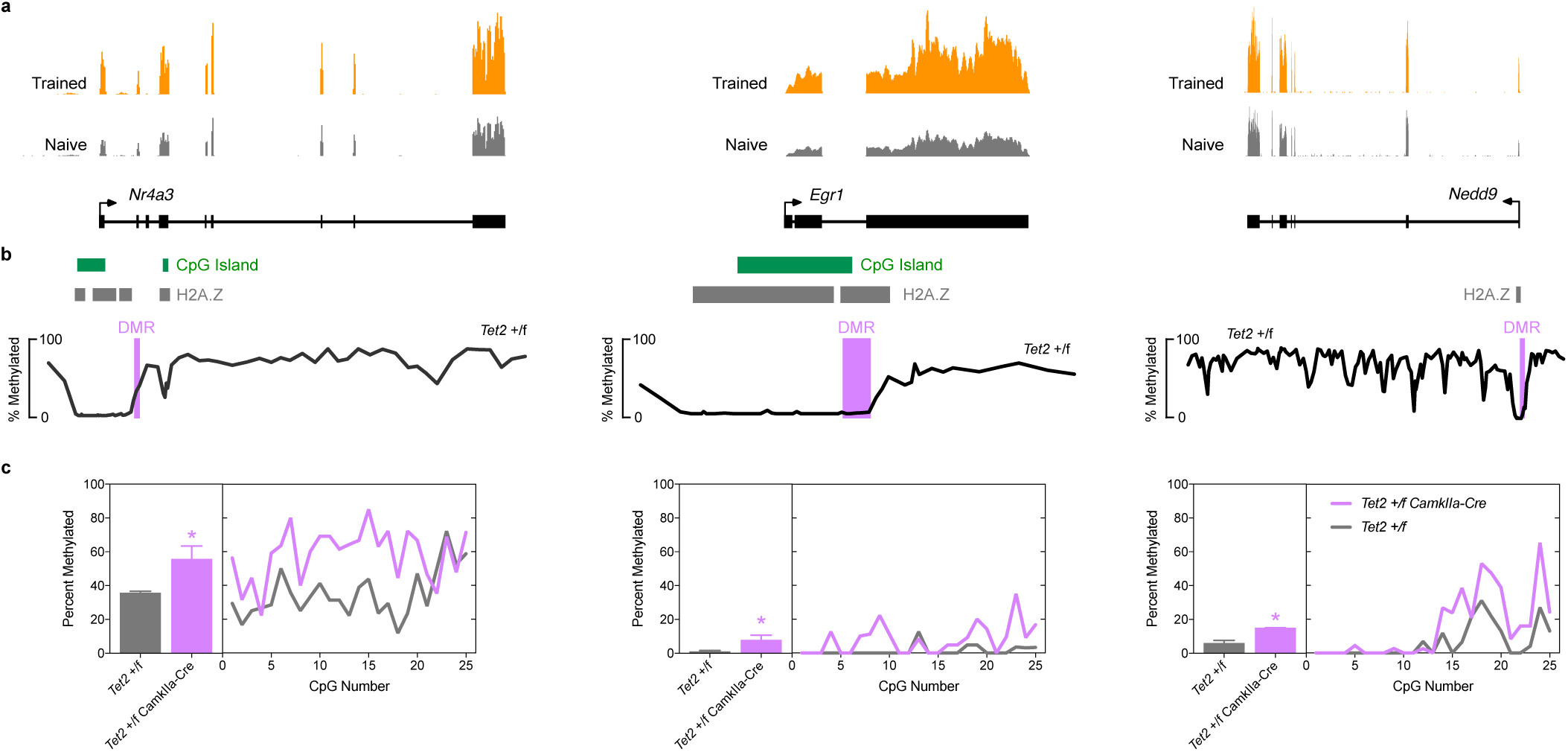
The conditional knockout of *Tet2* in glutamatergic neurons of the CA1 results in the hypermethylation of genes associated with memory formation. (A) Representative experience-regulated genes (*Nr4a3, Egr1*, and *Nedd9*) have significantly increased transcription 1 h following contextual fear conditioning as compared with naïve littermates. Poly A+ RNAseq data from the hippocampus of mice 1 h after contextual fear conditioning training (orange) or from naïve mice (grey). N = 6 per group. (B) DMRs are associated with Unmethylated regions around the TSS of these active genes. (C) The percent change in DNA methylation across the indicated DMR and the level of individual CpG methylation with in that DMR. Data shown are represented as mean ± sem. EdgeR filter for differential methylation in WGBS (FDR < 0.05) or DESeq2 filter for differential expression analysis in RNAseq (FDR < 0.05)

## Discussion

We interpret the data from this study to indicate that *Tet2* functions to de-methylate DNA at the promoters and TSSs of genes that are expressed after learning, which is associated with an increased fidelity in memory recall of those experiences. The deficits in memory extinction caused by *Tet2* reduction in glutamatergic circuits of the CA1 and forebrain indicate that *Tet2* activity does not modulate cognition generally, but selectively attenuates the strength of memory. Importantly, this study measures the modified DNA methylation that is the product of *Tet2* reduction in these circuits with single cytosine resolution using WGBS, which allows for the precise interrogation of the resulting hypermethylated loci. Whereas *Tet1* and *Tet3* have also been previously implicated in memory function and glutamatergic signaling, this study relates the activity of *Tet2* directly to a family of genes activated by experiential learning.

*Tet2* has recently been identified as a therapeutic target for Alzheimer’s Disease (Carrillo-Jimenez et al., 2019), and the data presented here suggest that *Tet2* inhibition in the CA1 of the hippocampus and in post-mitotic glutamatergic neurons is sufficient to enhance memory fidelity. Important previous work has shown that *Tet2* also regulates neurogenesis in the dentate gyrus of mice (Gontier et al., 2018). Being an epigenetic modifier, *Tet2* is necessary for proper neural progenitor cell (NPC) differentiation in the dentate gyrus, and the sustained knockout of *Tet2* in NPCs over the course of two months was demonstrated to impair cognitive function. We propose that these data compliment the present study in two fundamental ways. First, *Tet2-*mediated DNA de-methylation drives cognitive flexibility through neurogenesis in the hippocampus and by facilitating memory adaptation and extinction when the subject is presented with new information. Second, whereas complete ablation of *Tet2* in the CNS has long-term consequences for neurogenesis and cognition, therapeutic intervention that reduces, rather than eliminates, *Tet2*-mediated de-methylation of the genome in the hippocampus serves to enhance memory fidelity.

This study does not interrogate the effects of *Tet2* activity on glutamatergic electrophysiology or whether increased levels of *Tet2* activity is sufficient to impair memory recall. However, it clearly links *Tet2* function as a negative regulator of memory fidelity and to the DNA methylation status of genes important for long-term memory formation. Inhibitors of *Tet2* have recently been described (Chua et al., 2019), and this study also implicates *Tet2* as a target to enhance long-term memory therapeutically.

## Author Contributions

KEZ, CPG, HCS, and AJK designed, performed, and analyzed all experiments except the following. GAK performed rodent surgeries and expression analysis. MMC performed bioinformatic analyses. XZ, AHH, ARB, ARG, AR, and BGM performed or aided with animal behavior. CZ and HS designed the viral vectors.

## Acknowledgements

Research reported in this publication was supported by an Institutional Development Award (IDeA) from the National Institute of General Medical Sciences of the National Institutes of Health under grant number P20GM103423, the Pitt-Hopkins Research Foundation, the Orphan Disease Center’s 2019 MDBR Pilot Grant Program, the Sherman Fairchild Foundation, and Bates College. This work is dedicated to the memory of Kindal Kivisto.

## Methods

### Animals

All mice were acquired from The Jackson Laboratory, Bar Harbor, ME. Mice used for AAV injections were C57BL/6J mice (Stock # 000664), while *Tet2* flox (Stock # 017573) and Camk2a-Cre / T29 (Stock # 005359) mice were bred to produce *Tet2* +/flox Camk2a-Cre and *Tet2* +/flox littermates. All experiments were performed between 2 and 3 months of age. All procedures were performed with IACUC-approved protocols.

### Surgeries

ShRNA-containing AAVs were injected into the dorsal hippocampus (dHPC) using the stereotaxic coordinates, –2.0 mm AP, ±1.5 mm ML, and −1.6 mm DV relative to bregma. 1 uL of viral-containing solution was injected per hemisphere, as previously described (Kumar et al., 2015). Injections were performed using a 10 mL Hamilton Gastight syringe controlled by a Pump 11 Elite Nanomite Programmable Syringe Pump (Harvard Apparatus). Injections proceeded at a speed of 150 nL min−1 through a 32-gauge needle. The injection needle was left in place an additional 5 min. At 14 days post-surgery, EYFP-infected dHPC tissue was removed, snap frozen and stored at −80 °C until further use. Target sequences of shRNAs:

Tet2 shRNA target sequence

5’-GGATGTAAGTTTGCCAGAAGC-3’

Scrambled shRNA target sequence

5’-GTTCAGATGTGCGGCGAGT-3’

### Open Field Task

Anxiety-related behavior, thigmotaxis, and overall activity were assessed using the open field test. The open field apparatus is a 27 × 27 cm arena with 20 cm high walls on all sides. Animals were placed in the center of the field and allowed to explore the box for a single 30 min trial. The movement of animals was tracked using Activity Monitor software (Med Associates).

### Object Location Memory

The object location memory (OLM) task has been previously described to test spatial memory in rodents (Ciernia and Wood, 2014). The boxes used in OLM were made of white plastic with a 25.5 × 25.5 cm floor and 31 cm high walls. A blue piece of tape marked a center point on one of the four walls and mice were always placed in the center of the box facing the center mark to allow assurance of their location. Mice were handled and tested in a dimly lighted room with proportional amounts of light in each box (4-6 lux), as measured by an illuminance light meter (Sunche, HS1010). Training and testing trials were recorded with an overhead camera using EthoVision software (Noldus). Mice were handled for 4-6 days prior to the beginning of the OLM task. This consisted of a 1 m period of holding a mouse followed by a 5 m exploration period of a box that contained no objects. Training consisted of one 10 min exposure to two 50 mL beakers taped up-side down and equidistance from the blue cue. 2 h, 1 d, and 7 d following training, mice were tested on the OLM task to assess spatial memory. Testing consisted of a 5 m exploration interval in which one of the beakers was moved to a novel location (distal to the blue cue). All OLM training and testing videos taken by an overhead camera were scored by genotype blinded observers. The amount of time that mice spent interacting with each present object was measured and used to calculate a discrimination index (DI) that quantified the relative difference in time spent investigating the two objects. This index was defined as: DI = (N−F) / (N+F) x 100, where “N” was the amount of time spent investigating the object novel location and “F” was the amount of time spent investigating the object in the familiar location. Time spent investigating the object was considered to be any time that the animals spent with their noses to the object, excluding any time they spent sitting atop the object.

### Passive Avoidance

Animals were trained with a shuttle box (Med Associates) which consists of two chambers separated by a guillotine door. The leftmost chamber was always illuminated by a light while the rightmost chamber was covered with a dark cloth and left unlighted. Passive avoidance training began with the guillotine door down and animals were placed in the lighted chamber and allowed 20 s to habituate. The door was then opened, allowing for exploration of the dark chamber. Upon complete entrance to the dark chamber, the guillotine door closed, and animals were given a 2 s 0.5 mA foot shock. Animals were trained using either weak (1x training) or strong (3x training) fear training. For a weak training session, animals were immediately removed from the dark chamber following their first shock. After each shock in a strong training session, the guillotine door opened following a 30 s intertrial-interval in which animals were confined to the dark chamber. During testing, animals were placed in the lighted chamber, given a 20 s acclimation period before the guillotine door opened. Latency to reenter the dark chamber was recorded. Following weak passive avoidance training, mice were tested on the shuttle box 1 d after training. Mice given strong training were then submitted to 8 days of extinction testing. These animals underwent one test session per day in which they were allowed 5 m in the shuttle box to explore the dark chamber, and latency to reenter the dark chamber was measured.

### qRT-PCR method for Tet2 detection

Total RNA was extracted from AAV2-infected shRNA-scrambled and -TET2 dorsal hippocampi using an RNeasy Mini Plus Kit (Qiagen). 100 ng Total RNA was converted to cDNA using a iScript™ cDNA Synthesis Kit (Bio-Rad). qRT-PCR was performed on an CFX96 real-time PCR detection system in 10 *μ*L reactions containing SsoAdvanced™ Universal SYBR® Green Supermix and 200-300 *μ*M of primer and 1 *μ*L of 1:5 diluted cDNA in technical triplicates. Amplification of Tet2 cDNA transcripts were amplified using the pre-designed PrimeTime® qPCR Primer Assay Mm.PT.58.300089849 (IDT). Relative fold quantification of gene expression between samples was calculated using the delta-delta Ct method and normalized to the reference gene *Hprt1*.

### Whole Genome Bisulfite Sequencing

Hippocampi were extracted from 4-month-old mice (6 animals per genotype) and the CA1 region sub-dissected. gDNA was extracted (Qiagen, DNeasy), pooled into two samples per genotype (gDNA from 3 mice per sample), bisulfite converted (Accel-NGS Methyl-Seq DNA Library, Swift), and sequenced (Hudson Alpha Discovery) on the Illumina platform (HiSeq_X10, paired end, 150 bp, 500 million reads per sample). Reads were QC’d by FastQC (v0.11.5), filtered by quality (average quality score < 20), Illumina adapters removed, and sequencing primers trimmed using Trim_Galore! (v0.4.5) according to the Swift kit specifications. The filtered and trimmed reads were then mapped to the mouse genome (mm10), deduplicated, and CpG methylation levels extracted using Bismark (v0.19.0). Altered CpG methylation was determined by dividing the mouse genome into 849,172 windows, each containing 25 CpG sites. Differentially methylated regions (DMRs) were determined using the EdgeR (for/rev) algorithm available in the Seqmonk (v1.44.0) software package, where significant differences between genotypes were determined by an FDR < 0.05. H2A.Z binding sites were determined using previously published datasets (Stefanelli et al., 2018), as well as for determining active hippocampal promoters and enhancers (Gjoneska et al., 2015). Prior to bisulfite conversion, samples were spiked with unmethylated Lambda DNA. Reads were also mapped using Bismark to the Lambda genome to confirm cytosine hydrolysis. The fastq files for these datasets have been made available on GEO under the accession number GSE140575.

