## Supplemental Information for "*Tet2* negatively regulates memory fidelity"

**Table of Contents**

Figure S1 – *Tet2* Knockdown Baseline Behavior ........................................................................2
Figure S2 – *Tet2* Knockout Baseline Behavior ..........................................................................3
Table S1 – Annotated DMRs ........................................................................Attached text file

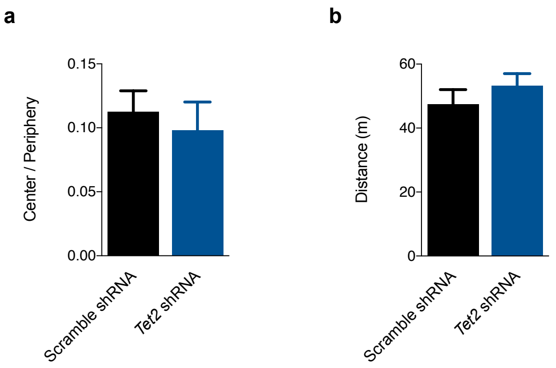


**Figure S1.** ***Tet2*** **Knockdown Baseline Behavior, related to Figure 1.**

(A) *Tet2*-shRNA mediated knockdown in CA1 did not significantly affect thigmotaxis over a 30-minute open field test (N = 7 per group).

(B) *Tet2*-shRNA knockdown did not significantly affect ambulation in the open field.

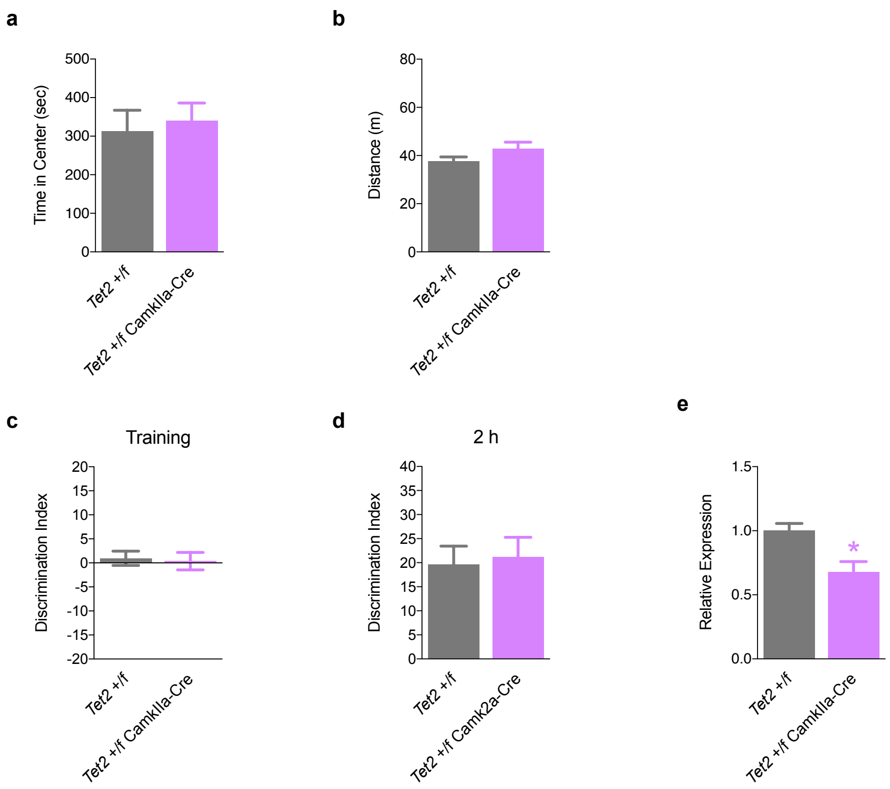


**Figure S2.** ***Tet2*** **Knockout Baseline Behavior, related to Figure 2.**

(A) *Tet2* +/flox Camk2a-Cre mice did not elicit significantly altered thigmotaxis over a 30-minute open field test as compared with *Tet2* +/flox mice (N = 17-22 per group).

(B) *Tet2* +/flox Camk2a-Cre mice did not demonstrate significantly affect ambulation in the open field as compared with *Tet2* +/flox mice in the open field (N = 17-22 per group).

(C) Neither *Tet2* +/flox Camk2a-Cre or *Tet2* +/flox mice showed any preference for the objects during OLM training (N = 27-28).

(D) *Tet2* +/flox Camk2a-Cre mice did not show altered short-term memory at 2 hours after training in OLM, as compared with *Tet2* +/flox mice (N = 13-14 per group)

(E) Relative levels of *Tet2* mRNA determined by from the CA1 region of the hippocampus in *Tet2* +/flox Camk2a-Cre and *Tet2* +/flox mice (N = 5 per group)
